# Reading-related brain changes in audiovisual processing: cross-sectional and longitudinal MEG evidence

**DOI:** 10.1101/2021.05.19.444836

**Authors:** Sendy Caffarra, Mikel Lizarazu, Nicola Molinaro, Manuel Carreiras

## Abstract

The ability to establish associations between visual objects and speech sounds is essential for human reading. Understanding the neural adjustments required for acquisition of these arbitrary audiovisual associations can shed light on fundamental reading mechanisms and help reveal how literacy builds on pre-existing brain circuits. To address these questions, the present longitudinal and cross-sectional MEG studies characterize the temporal and spatial neural correlates of audiovisual syllable congruency in children (4-9 years old, 22 males and 20 females) learning to read. Both studies showed that during the first years of reading instruction children gradually set up audiovisual correspondences between letters and speech sounds, which can be detected within the first 400 ms of a bimodal presentation and recruit the superior portions of the left temporal cortex. These findings suggest that children progressively change the way they treat audiovisual syllables as a function of their reading experience. This reading-specific brain plasticity implies (partial) recruitment of pre-existing brain circuits for audiovisual analysis.

**Significance Statement:** Linking visual and auditory linguistic representations is the basis for the development of efficient reading, while dysfunctional audiovisual letter processing predicts future reading disorders. Our developmental MEG project included a longitudinal and a cross-sectional study; both studies showed that children’s audiovisual brain circuits progressively change as a function of reading experience. They also revealed an exceptional degree of neuroplasticity in audiovisual neural networks, showing that as children develop literacy, the brain progressively adapts so as to better detect new correspondences between letters and speech sounds.

## 1 Introduction

Literacy is a relatively recent cognitive achievement in human evolution for which there are no specialized neural circuits already in place. Learning this life-changing skill thus requires considerable modulation of pre-existing brain networks, such as the visual object recognition and spoken language networks (Carreiras et al., 2009; Dehaene, Cohen, Morais, & Kolinsky, 2015). A considerable amount of research on reading-related brain changes has examined this plasticity in either visual and auditory brain circuits (Dehaene et al., 2010, 2015; Goswami & Ziegler, 2006; Ziegler & Muneaux, 2007). However, the core of reading acquisition lies in the interaction between these two modalities. Efficient reading skills crucially depend on the ability to compare and connect visual and auditory representations of letters (Blomert, 2011). The present MEG study focused on these audiovisual processes, testing how they changed as a function of developing reading abilities. We hypothesized that during reading acquisition pre-existing brain circuits for audiovisual processing should become progressively tuned to the arbitrary relationships between letters and speech sounds (Blomert, 2011).

The processing of natural audiovisual associations (e.g., the correspondence between speech and lip movements) has been widely explored in the literature. The effects of audiovisual integration (i.e., the absolute difference between bimodal and unimodal presentations) and audiovisual congruency (i.e., the absolute difference between matching and mismatching bimodal presentations) have mainly been localized in the auditory cortex and the superior temporal cortex (Amedi, von Kriegstein, van Atteveldt, Beauchamp, & Naumer, 2005; Hocking & Price 2008), with possible left lateralization (Calvert, 2001; Calvert, Brammer, & Iversen, 1998). Research on fluent adult readers has shown that these brain areas seem to be (at least partially) recruited even in processing arbitrary associations between letters and speech sounds, indicating a certain degree of plasticity in audiovisual brain areas during reading acquisition (Amedi et al., 2005; Blomert & Froyen, 2010; Hocking & Price, 2008). Neuroimaging studies comparing matching and mismatching letter-sound pairs reported effects in the superior temporal and auditory cortex (Blau, van Atteveldt, Formisano, Goebel, & Blomert, 2008; Blau, Reithler, van Atteveldt, Seitz, Gerretsen, Goebel, & Blomert, 2010; Karipidis et al., 2017, 2018; van Atteveldt, Formisano, Goebel, & Blomert, 2004; van Atteveldt, Formisano, Blomert, & Goebel, 2007), which were often left-lateralized and appeared within the first 500 ms of stimulus presentation (Herdman, Fujioka, Chau, Ross, Pantev, & Picton, 2006; Karipidis et al., 2017, 2018; Raji, Uutela, & Hari, 2000; Xu, Kolozsvári, Oostenveld, Leppänen, & Hämäläinen, 2019; Xu, Kolozsvári, Oostenveld, & Hämäläinen, 2020; for even earlier effects see Herdman et al., 2006). Importantly, cross-sectional designs have revealed a relation between these audiovisual effects and reading skills (Blau et al., 2010; Karipidis et al., 2017, 2018; cfr. Jost, Eberhard-Moscicka, Frisch, Dellwo, & Maurer, 2014), indicating that cross-modal brain responses are affected by literacy experience. Studies on normal reading acquisition in children seem to suggest that automatic effects of audiovisual letter processing are rare in beginning readers (Xu, Kolozsvari, Monto, & Hämäläinen, 2018) and may emerge only after a few years of formal reading instruction under facilitated experimental conditions (e.g., non-simultaneous bimodal presentations, Froyen, Bonte, van Atteveldt, & Blomert, 2009). However, the scarce research on these plastic brain changes during development has so far been documented only by means of between-group comparisons. Longitudinal designs overcome the potential limitations – related to the difficulty of establishing perfectly matching groups – in between-group designs. The present MEG study is the first to adopt a longitudinal (alongside a cross-sectional) design in order to characterize the progressive emergence of audiovisual congruency effects as children learn to read. Matching and mismatching audiovisual syllables were presented to children. We predicted that the audiovisual congruency effect should be localized in the left superior temporal cortex and left auditory cortex and emerge within 500 ms after stimulus onset. We expected this effect to be reading-specific and, thus, to correlate with children’s reading scores.

## 2 Materials and Methods

### 2.1 Participants

Forty-two Basque-Spanish early bilingual children participated in the cross-sectional study (20 females, mean age: 6.3 years, SD: 1.7, age range: 4-9). Data from five additional participants were excluded due to poor data quality (n=4) or the presence of a hearing disorder (n=1). Participants were divided in two groups (pre-readers and readers) based on whether they had already received formal reading instruction (see Table 1). Fifteen children from the pre-readers group also participated in the longitudinal study, returning for a second MEG recording session. The mean time between Session 1 and Session 2 was 32 months (SD: 5, age range: 4-8, see Table 1).

**Table 1.** Behavioral description of participants in the cross-sectional and the longitudinal studies. For each measure, we included mean, standard deviation, and range of raw scores. The nonverbal intelligence quotient (IQ) was assessed using the Raven’s progressive matrices test and vocabulary size was measured using a picture naming task (both taken from Kaufman brief intelligence test, K-BIT, Spanish version, 2009). Reading performance was assessed by measuring speed and number of errors on 30 Basque syllables (the MEG experimental stimuli), 80 Basque high-frequency words and 80 Basque pseudowords.

|  | Cross-sectional study |  | Longitudinal study |  |
| --- | --- | --- | --- | --- |
|  | Pre-readers<br>M(SD) | Readers<br>M(SD) | Session 1<br>M(SD) | Session 2<br>M(SD) |
| Nº | 20 | 22 | 15 | 15 |
| Nº females | 9 | 11 | 7 | 7 |
| Nº right-handed | 12* | 21 | 8* | 12 |
| Age - years | 4.8 (0.8) | 7.7 (0.7) | 4.5 (0.7) | 7.1 (0.3) |
| Formal reading instruction duration-months | 0 | 14.2 (4.7) | 0 | 8.1 (1.7) |
| Nonverbal IQ (0-48) | 14.1 (5.3) | 25.6 (4.2) | 12.3 (4.6) | 24.8 (3.0) |
|  | 5-24 | 15-34 | 5-22 | 20-30 |
| Vocabulary size (0-45) | 20.4 (5.1) | 32.9 (4.2) | 18.9 (4.4) | 30.3 (4.2) |
|  | 12-31 | 23-38 | 12-28 | 23-38 |
| Letter knowledge (0-27) | 13.7 (10.0) | 27(0.0) | 10.8 (8.6) | 27(0.0) |
|  | 0-27 | 27-27 | 0-27 | 27-27 |
| Syllables reading times - sec | - | 21.6 (6.9) | - | 26.7 (8.7) |
|  | - | 14-43 | - | 17-46 |
| Word reading errors (0-80) | - | 2.4 (2.4) | - | 4.6 (4.0) |
|  | - | 0-9 | - | 0-15 |
| Word reading times - sec | - | 85.1 (42.2) | - | 133.7 (54.7) |
|  | - | 41-248 | - | 62-245 |
| Pseudoword reading errors (0-80) | - | 3.3 (1.9) | - | 6.6 (5.5) |
|  | - | 1-8 | - | 0-20 |
| Pseudoword reading times - sec | - | 90.5 (42.6) | - | 125.9 (49.6) |
|  | - | 42-236 | - | 61-245 |

\*The rest of the children did not show a clear hand preference at this time point.

All participants were learning to read in Basque. Basque has a transparent orthography, such that the consistent correspondences between letter and speech sounds are usually mastered within one year of reading instruction. Readers’ school attendance was regular and none of them were repeating or had skipped a grade. All participants had normal or corrected-to-normal vision, normal hearing. Their parents reported no neurological disorders and did not suspect developmental reading problems. The BCBL ethical committee approved the experiment (following the principles of the Declaration of Helsinki) and all parents (or the tutors) of the children compiled and signed the written informed consent.

### 2.2 Materials and procedure

Thirty consonant-vowel syllables were created using one of 6 consonants (f, k, l, m, p, t) followed by one of 5 vowels (a, e, i, o, u) from the Basque alphabet. We used syllables rather than single letters to make the stimuli more ecological. Basque children learn to name Basque letters using syllables and the consonant-vowel syllable structure is highly common in the Basque lexicon. We did not expect this choice to affect our results as audiovisual congruency effects have been reported for a wide range of linguistic (e.g., letters, words, ideograms; Amedi et al. 2005, Hocking & Price, 2008; Xu et al., 2019) and non-linguistic (pictures; Hocking & Price, 2008) stimuli. The syllables were presented four times both in the visual and the auditory modality to create 120 cross-modal pairs. Spoken syllables were recorded by a female voice at 44.1 KHz. The audiovisual correspondence of cross-modal pairs was manipulated to produce 60 matching and 60 mismatching pairs. The mismatching pairs were pseudo-randomly selected so that they always differed in the initial consonant while sharing the final vowel. Sixteen cross-modal syllable pairs were added for a target detection task. They contained the image of a cat in between the letters in the visual presentation and/or the sound of a cat meowing in between the letter sounds in the auditory presentation.

During the experimental trial, the visual stimulus (written syllable) was first presented at the center of the screen. After a one second, the auditory stimulus (spoken syllable) was also presented, while the written syllable remained displayed on the screen. The visual and auditory stimuli offsets coincided and the interstimulus interval was 1000 ms (see Figure 1). The onsets of the visual and auditory syllable presentations were shifted in order to create a facilitated experimental situation where it was more likely to observe early audiovisual congruency effects (Froyen et al., 2009). Moreover, this temporal sequence better reflected children’s everyday experience, such as listening to stories read aloud, where they hear language after seeing it in print. Auditory stimuli were presented between 70 and 80 dB through plastic tubes and silicon earpieces (mean duration: 700 ms, SD: 95). The task consisted of pressing a button whenever the current stimulus corresponded to a cat either in the visual or in the auditory modality. Stimuli were randomized across participants. The recording session lasted approximately 10 minutes.

**Figure 1.**
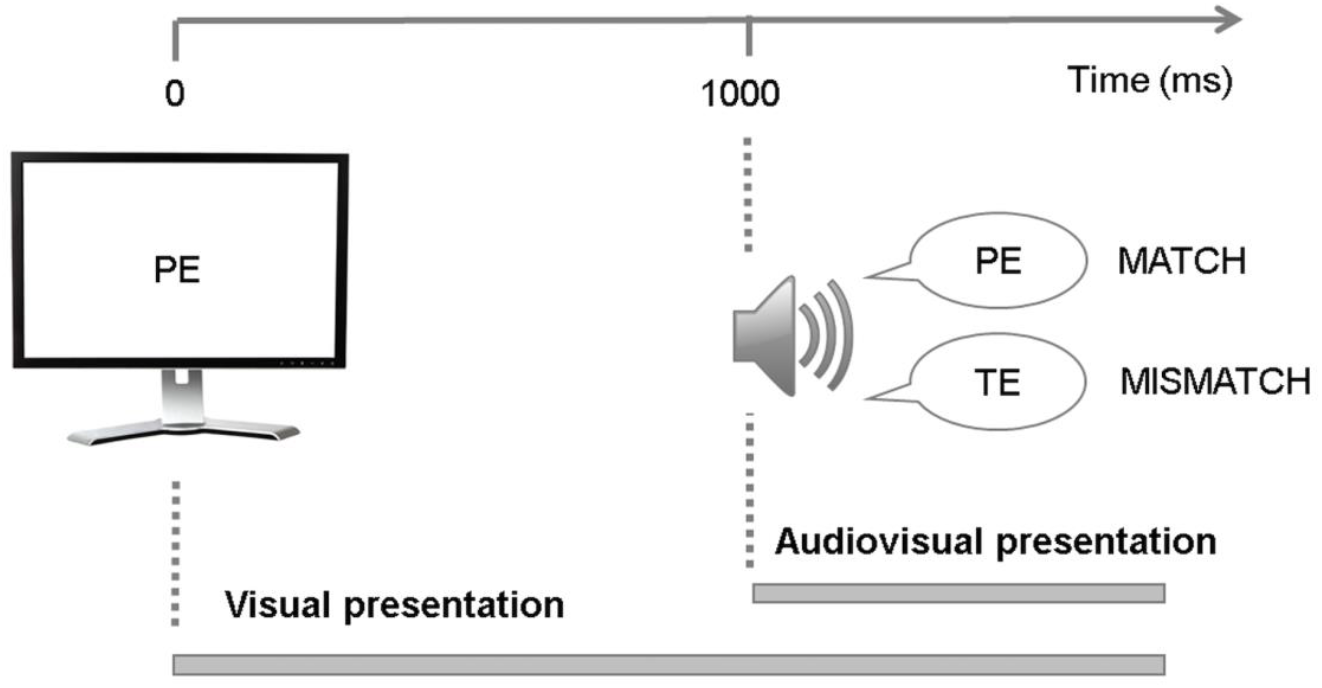
Schematic representation of an experimental trial.

### 2.4 MEG data recording and preprocessing

MEG data were recorded in a magnetically shielded room (Maxshieldł, Elekta Oy, Helsinki, Finland) using an Elekta-Neuromag MEG device (including 102 sensors with two planar gradiometers and one magnetometer each). MEG recordings were acquired continuously with children in a sitting position, with a bandpass filter at 0.03−330 Hz and a sampling rate of 1 KHz. Head position inside the helmet was continuously monitored using five head position indicator coils. The location of each coil relative to the anatomical fiducials (nasion, and left and right preauricular points) was defined with a 3D digitizer (Polhemus Fastrak, Colchester, VT, USA). This procedure is critical for head movement compensation during the data recording session. In addition, about 200 head surface points were digitized and later used to spatially align the MEG sensors with an age-based pediatric T1 template (Fonov, Evans, Botteron, Almli, McKinstry & Collins, 2011).

Eye movements were monitored with vertical and horizontal bipolar electrooculograms (VEOG and HEOG). MEG data were individually corrected for head movements and subjected to noise reduction using MaxFilter (Ver.2.2.15; Elekta-Neuromag) and the temporally extended signal space separation method (Taulu & Hari, 2009; Taulu & Kajola, 2005). On average, ten bad channels were automatically identified using Xscan (Elekta-Neuromag). Bad channels were substituted with interpolated values. There was no difference between the number of channels interpolated between readers (10.2, SD: 2.2) and pre-readers (9.1, SD: 2.2; *t*<1), or between Session 1 (10.1, SD: 2.3) and Session 2 (10.2, SD: 3.3; *t*<1).

Subsequent analyses were performed using Matlab R2014 (MathworksR©, Natick, MA) and the Fieldtrip toolbox (Oostenveld, Fries, Maris, & Schoffelen, 2011). MEG epochs of 2.5 seconds were obtained, including 1.5 sec before and 1.0 sec after the auditory presentation onset. High-frequency muscle artifacts (110-140 Hz) were automatically rejected: average z-values over sensors and time points in each trial were calculated and trials exceeding the threshold of a z-score equal to 30 were removed. To suppress eye-movement artifacts, 70 independent components were identified by applying Independent Component Analysis (ICA; Jung, Makeig, Humphries, Lee, McKeown, Iragui, & Sejnowski, 2000) to the MEG data. Independent components corresponding to ocular artifacts were identified and removed based on the correlation values between each component and the VEOG/HEOG channels (rejected components range: 0-2).

Finally, MEG epochs were visually inspected to discard any remaining artifacts. On average, 28.1% (SD: 13.1) of trials were rejected (cross-sectional study: 26.7%, 11.6; longitudinal study: 30.0%, 14.9), with no significant difference between conditions (*F*s<3, *p*s>.05) or groups (*F*s<5, *p*s>.05).

### 2.5 MEG experimental design and statistical analysis

*Sensor-level ERFs:* The artifact-free MEG data were lowpass filtered below 35 Hz. Trials were grouped together for each condition and then averaged to obtain the Event Related Fields (ERFs). ERFs were quantified as the absolute amplitude of the 102 orthogonal planar gradiometer pairs by computing the square root of the sum of squares of the amplitudes of the two gradiometers in each pair. A baseline correction for the data preceding the stimulus by 500 ms was performed.

In both the cross-sectional and longitudinal studies, the ERFs for the match and mismatch conditions of pre-readers and readers were statistically compared using a nonparametric cluster-based permutation test (Maris & Oostenveld, 2007). Specifically, *t*-statistics were computed for each sensor (combined gradiometers) and time point during the [0 – 1000] ms time window, and a clustering algorithm formed groups of channels over time points based on these tests. The neighborhood definition was based on the template for combined gradiometers of the Neuromag-306 provided by the toolbox. In order for a data point to become part of a cluster, a threshold of p = 0.05 was used (based on a two-tailed dependent t-test, using probability correction). The sum of the t-statistics in a sensor group was then used as a cluster-level statistic (e.g., the maxsum option in Fieldtrip), which was then tested with a randomization test using 1000 runs. Moreover, we used a two tailed t-test to perform a between-group comparison of the audiovisual congruency effects (ERF differences between mismatch and match conditions) in the cross-sectional and the longitudinal study. Finally, partial correlations were calculated to evaluate the relationship between the magnitude of the audiovisual congruency effect and reading performance after correcting for age, vocabulary size, and nonverbal intelligence.

*Source-level ERFs*: Using MRiLab (Elekta Neuromag Oy, version 1.7.25), the digitized points from the Polhemus were co-registered to the skin surface obtained from an age-compatible T1 template (Fonov et al., 2011). The T1 template was segmented into scalp, skull, and brain components using the segmentation algorithms implemented in Freesurfer (Martinos Center of Biomedical Imaging, MQ). The source space was defined as a regular 3D grid with a 5 mm resolution and the lead fields were performed using a realistic three-shell model. Both planar gradiometers and magnetometers were used for inverse modelling. Whole brain source activity was estimated using the linearly constrained minimum variance (LCMV) beamformer approach (Van Veen, van Drongelen, Yuchtman, & Suzuki, 1997). For each condition, LCMV beamformer was computed on the evoked data in the -400 to 0 pre-stimulus and in the 350 to 750 msec post-stimulus time intervals. This post-stimulus interval was chosen because it contained the audiovisual congruency effects at the sensor level. Statistical significance was assessed by a paired t-test (from Statistical Parametric Mapping software) comparing mean amplitudes in the post and the pre-stimulus interval (SPM).

## 3 Results

Participants were able to correctly identify the target stimuli (cross-sectional d’: 1.870; longitudinal d’: 2.077), with no differences across groups (cross-sectional: *t*(40)= 1.114, *p* = 0.272; longitudinal: *t*(14)= 1.872, *p*=0.082).

For the cross-sectional study (Figure 2A), cluster-based permutations on the ERF responses showed an audiovisual congruency effect (p=0.001) (difference between mismatch and match condition) only for readers in a 350–790 ms time window following the auditory syllable onset over left temporal sensors (highlighted in Figure 2A in the top left corner). The magnitude of the audiovisual congruency effect differed between readers and pre-readers (*p*= 0.005). This difference was due to the suppressed amplitude of the match condition in readers compared to pre-readers (match condition: *p*=0.021; mismatch condition: p= 0.105; Figure 2B).

**Figure 2.**
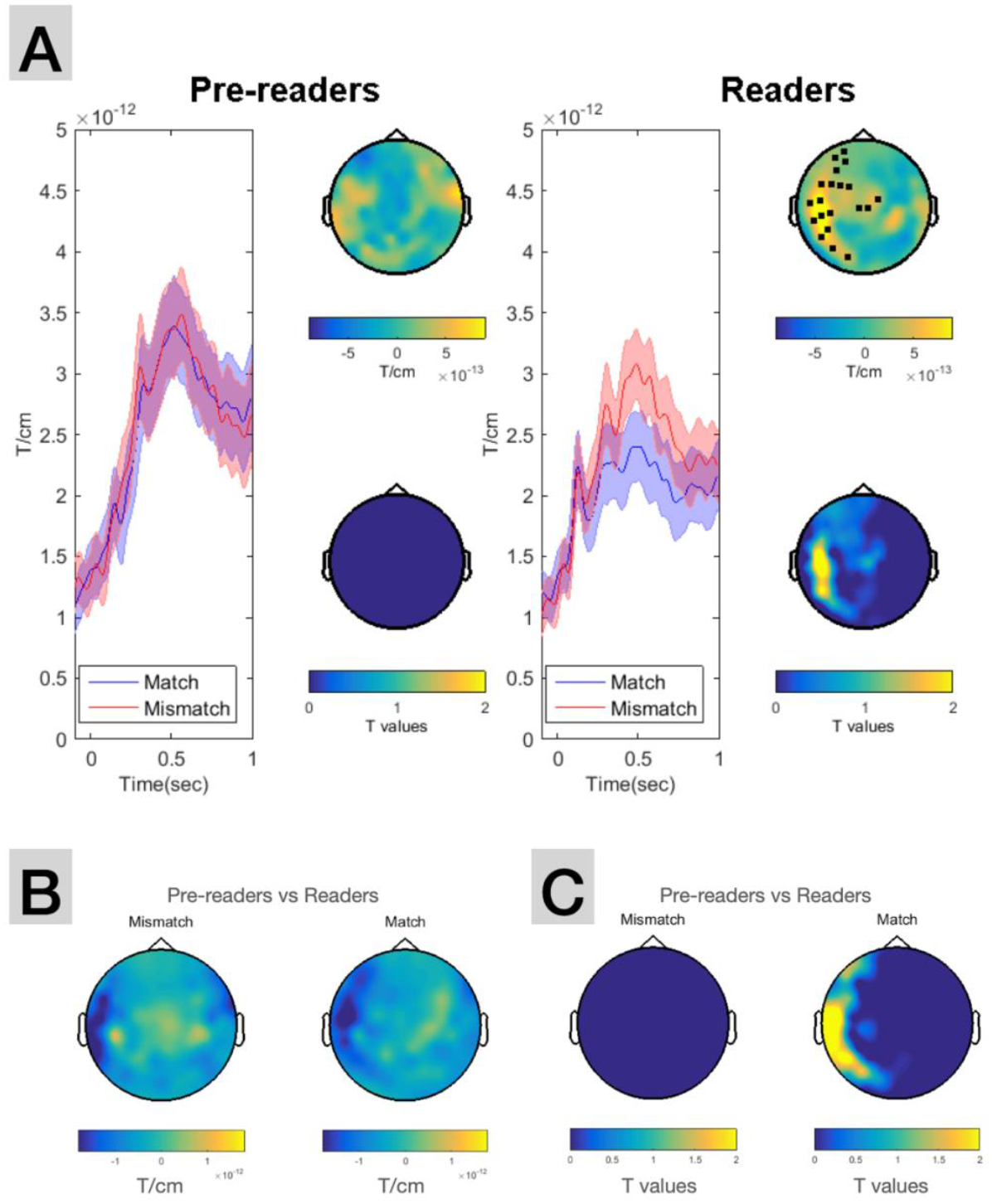
A: ERFs for the cross-sectional study. A: Grand average ERF responses to spoken syllables for the match (blue) and the mismatch (red) condition in pre-readers (left panel) and readers (right panel). Shaded edges represent +/-1 standard error. ERF waveform averages were calculated based on the group of left sensors displayed on the map in the left upper corner. The top maps represent the topographic distribution of the audiovisual congruency effect (calculated by subtracting the match from the mismatch condition) within the time window when the effect reached its maximum. The topographic maps at the bottom show the spatial distribution of the statistically significant cluster in the same time window (yellow color scale indexes the magnitude of *t* values that passed the statistical threshold of 0.05). B: Topographic maps of the difference between readers and pre-readers. C: Spatial distribution of the statistically significant cluster when comparing readers and pre-readers (yellow color scale indexes significant t values magnitude).

Similarly, for the longitudinal study (Figure 3A), we observed an audiovisual congruency effect (p=0.017) in a 390–563 ms time window following the auditory syllable onset over left temporal sensors (highlighted in Figure 3A top left corner). The magnitude of the audiovisual congruency effect differed between sessions (Session 1 vs Session 2: *p*= 0.038). Again, this difference was due to the suppressed amplitude of the match condition in the readers (Session 2) as compared to the pre-readers (Session 1, match condition: *p*=0.021; mismatch condition: p=0.627; Figure 3B).

**Figure 3.**
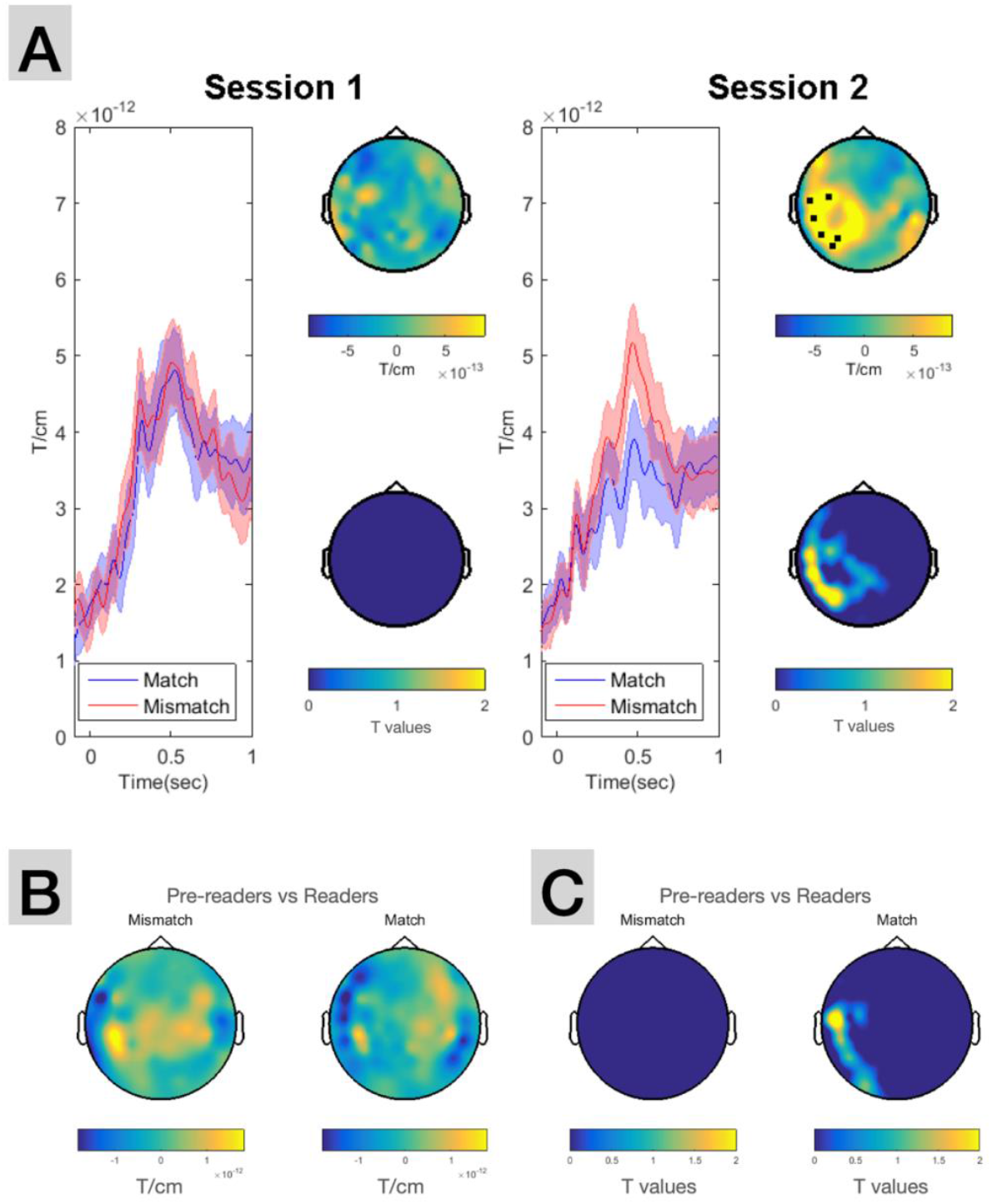
A: ERFs for the longitudinal study. Grand average ERF responses to spoken syllables for the match (blue) and the mismatch (red) condition in Session 1 and Session 2. Shaded edges represent +/-1 standard error. ERF waveform averages were calculated based on the group of left sensors displayed on the map in the left upper corner. The top maps represent the topographic distribution of the audiovisual congruency effect (calculated by subtracting the match from the mismatch condition) within the time window when the effect reached its maximum. The topographic maps at the bottom show the spatial distribution of the statistically significant cluster in the same time window (yellow color scale indexes the magnitude of *t* values that passed the statistical threshold of 0.05). B: Topographic maps of the difference between Session 1 and Session 2. C: Spatial distribution of the statistically significant cluster when comparing Session 1 and Session 2 (yellow color scale indexes significant t values magnitude).

The ERF effects observed at the sensor level were source reconstructed in the 350 to 750 ms time window. In both the cross-sectional and longitudinal study the congruency effect (p<0.05) emerged in the posterior part of the left superior temporal cortex (Figure 4).

**Figure 4.**
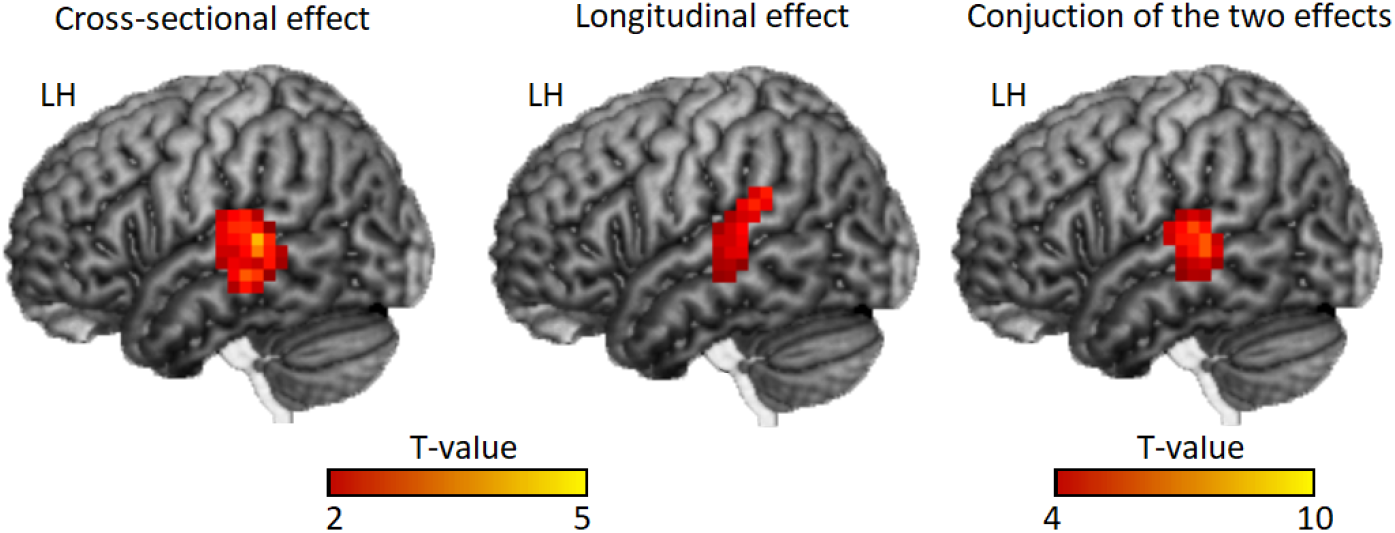
Spatial localization of the audiovisual congruency effect for readers of the cross-sectional and the longitudinal study. Paired t-test comparing the mean source activity in the pre-and post-stimulus interval were compared. Different color intensity indexes significant *t* values.

The size of the audiovisual congruency effect negatively correlated with reading errors and reading speed measures after correcting for age, non-verbal intelligence, and vocabulary size (syllable reading times: r= -0.31, p= 0.031; number of errors per second while reading Basque words: r= -0.36, p= 0.014; number of errors per second while reading Basque pseudowords: r= -0.23, p= 0.090; see Figure 5).

**Figure 5.**
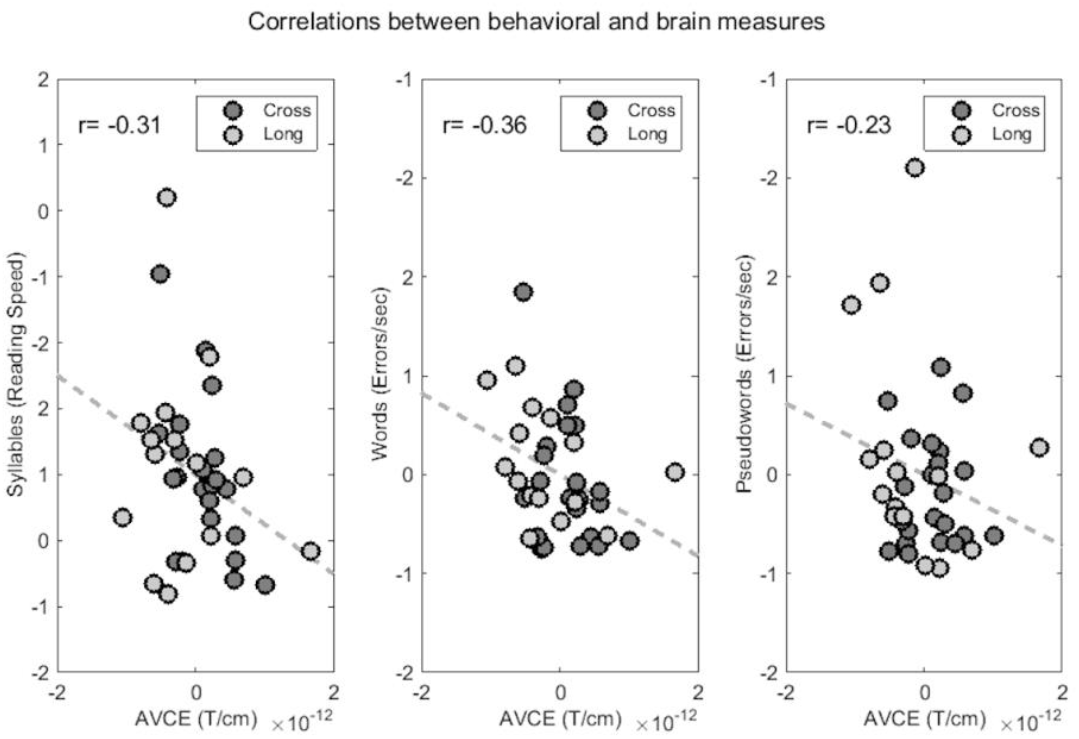
Correlation between the residuals of the audiovisual congruency effect (AVCE) and the residuals of reading scores (after correction for age, non-verbal intelligence, and vocabulary size). From left to right: syllable reading times, number of errors per second while reading Basque words, number of errors per second while reading Basque pseudowords. All readers are displayed in the scatterplots (*n*= 37; dark gray: cross-sectional study; light gray: longitudinal study).

## 4 Discussion

The capacity to create strong associations between speech sounds and written representations is a key skill for reading. Audiovisual letter and audiovisual symbol processing predict future reading fluency (Horbach, Scharke, Cröll, Heim & Günther, 2015; Horbach et al., 2018; Karipidis et al., 2018) and are often impaired in dyslexia (Fox, 1994; Froyen, Willems & Blomert, 2011; Richlan, 2019; Vellutino, Fletcher, Snowling, & Scanlon, 2004). Understanding the developmental changes involved in letter to speech sound processing can shed light on the pivotal mechanisms of reading and can point to possible sources of reading disorders. With this aim, the present study investigated how audiovisual syllable analysis changed as a function of reading acquisition. The results showed a high degree of plasticity in neural responses to audiovisual syllable congruency, which was related to reading acquisition (as shown by partial correlations with reading performance). This neural adjustment was mainly localized in the left superior temporal cortex, in line with previous findings (Blau et al., 2008, 2010; Karipidis et al., 2017, 2018; Raji et al., 2010; Xu et al., 2019, 2020). Importantly, this brain area is not exclusively involved in the processing of letter-speech sound correspondences, but also sensitive to less arbitrary audiovisual associations available before reading acquisition (Amedi et al., 2005; Calvert et al., 1998; Calvert, 2001). This broad sensitivity is compatible with the idea that we do not have evolutionarily specialized circuits for reading, and literacy must build on pre-existing brain networks (Dehaene et al., 2010, 2015). In line with this hypothesis, previous findings have shown reading-related adjustment of naturally evolved brain mechanisms for visual and auditory processing (Dehaene et al., 2010, 2015; Goswami & Ziegler, 2006; Ziegler & Muneaux, 2007). The present findings extend this claim, suggesting that reading experience can also have an impact on naturally evolved brain mechanisms for audiovisual processing (Blomert, 2011).

The direction of the audiovisual congruency effect is also informative. Past research reveals considerable inconsistency: some studies have shown stronger responses for matching conditions; others report the opposite pattern (see Table 2). Although it remains unclear what drives the direction of the effect (some proposals can be found in Hollaway, van Atteveldt, Blomert, & Ansari, 2018; Hocking & Price, 2008; Plewko et al., 2018; Wang, Karipidis, Pleisch, Fraga-Gonzalez, & Brem, 2020), we note that around 70% of the studies reporting stronger matching responses are fMRI studies. The reverse pattern has been more frequently observed in electrophysiology and with experimental designs that include non-simultaneous audiovisual presentations. This could indicate that temporal aspects of experimental design may affect the direction of the effect. The present MEG studies fully align with these trends found in the literature.

**Table 2.** Quick summary of the direction of audiovisual congruency effects (AVCE) previously reported in the literature. SOA: Stimulus-onset asynchrony. A summary of the timing and the MRI coordinates of the AVCEs is also reported in the two last columns (whenever available). aSTP: anterior superior temporal plane. HS: Heschl’s sulcus. IFG: inferior frontal gyrus. ITG: inferior temporal gyrus. MFG: middle frontal gyrus. MTC: middle temporal cortex. PT: planum temporale. STC: superior temporal cortex. STS: superior temporal sulcus.

|  | Technique | Participants | SOA for AVCE (ms) | Location (Time, ms) | Location (Space) |
| --- | --- | --- | --- | --- | --- |
| **Match > mismatch** |  |  |  |  |  |
| Van Atteveldt et al. 2004 | fMRI | adults | 0 | - | Bilateral PT/HS |
| Van Atteveldt et al. 2007 | fMRI | adults | Only at 0 | - | Bilateral PT/aSTP |
| Blau et al. 2009 | fMRI | children (~9 y) | 0 | - | Left STS |
| Blau et al. 2010 | fMRI | adults | 0 | - | Bilateral STS/Left PT/HS |
| Plewko et al. 2018 | fMRI | children with risk for dyslexia (~7y) | 0 | - | Left PT/STG |
| Karipidis et al. 2018 | fMRI | Pre-readers (~6y, future normal readers) | 0 | - | Left PT |
| Wang et al. 2020 | fMRI | children (poor readers, ~8y) | 0 | - | Right MFG |
| Raji et al. 2000 | MEG | adults | 0 | 415-515 | Bilateral STS |
| Herdman et al. 2006 | MEG | adults | 0 | 0-250 | Left PT/HS |
| Xu et al. 2020 | MEG | adults | 0 | 500-800 | Bilateral STC |
| **Mismatch > match** |  |  |  |  |  |
| Froyen et al. 2008 | EEG | adults | Max at 0 | ~160 | - |
| Froyen et al. 2009 | EEG | children (11y) | Only at 200 | ~160 | - |
| Froyen et al. 2011 | EEG | dyslexic children (11y) | Only at 200 | ~690 | - |
| Karipidis et al. 2017 | EEG/fMRI | Pre-readers (~6y) | SOA 0 | 382-442/644-704 | Right ITG |
| Karipidis et al. 2018 | EEG | Pre-readers (~6y, future normal readers) | SOA 0 | 382-442 | - |
| Xu et al. 2019 | MEG | adults | SOA 0 | 490-890 | Left STC/MTC |
| Hocking & Price, 2008 | fMRI | adults | Max at 0 | - | Left STS |
| Hollaway et al. 2015 | fMRI | adults | SOA 0 | - | Bilateral frontoparietal network |
| Plewko et al. 2018 | fMRI | children with no risk for dyslexia (7y) | SOA 0 | - | Left PT/STG |
| Wang et al. 2020 | fMRI | children (~8y, poor and normal readers) | SOA 0 | - | Right MTG/Left IFG/ITG/Bilateral SFG |

In both the longitudinal and cross-sectional study, we observed progressive suppression of the audiovisual matching response as a function of reading skills. Given that the congruency effect was found in auditory areas and the lack of modulation in the mismatch condition, it is unlikely that attention mechanisms accounted for this effect. This pattern is more likely the result of cross-modal integration since audiovisual correspondences can only be detected given successful interaction between two unimodal inputs. However, not all brain areas showing a congruency effect are necessarily the source of integrative operations (van Atteveldt et al., 2004, 2007; van Atteveldt & Anasari, 2014). Neuroimaging studies on adults comparing unimodal and bimodal letters proposed a finer functional distinction within subareas of the left superior temporal cortex. According to this view, the superior temporal sulcus is the neural hub for audiovisual convergence and integration, which sends feedback to superior auditory areas signalling letter-sound congruency (van Atteveldt et al., 2004, 2007). This functional distinction is further confirmed by cytoarchitectonic studies in human and non-human primates, which have shown a difference in the cellular structure of dorsolateral and ventromedial temporal regions (Ding, Hoesen, Cassell, & Poremba, 2009; Insausti, 2013; Zachlod et al. 2020). The reduced response of the superior temporal cortex to matching audiovisual syllables might reflect the sharpening of neuronal tuning (i.e., responses to overlearned audiovisual associations are suppressed; Hurlbert, 2000), cross-modal repetition suppression (Henson, 2003) or neural adaptation (Grill-Spector & Malach, 2001).

The present MEG results also support the idea that written letters systematically modulate children’s response to speech sounds in the left superior temporal cortex (Froyen et al., 2008; 2009; Herdman et al., 2006; van Atteveldt et al., 2007). Our longitudinal findings suggest that this effect is already present after a few months of formal reading instruction. A longer training period might be needed in order to reach a high degree of automaticity (and a shorter time window for audiovisual integration, Froyen et al., 2009; Laasonen, Tomma-Halme, Lahti-Nuuttila, Service, & Virsu, 2000; Laasonen, Service, & Virsu, 2002). In the present study, the long SOA between the visual and auditory onsets, together with the relatively late latency of our audiovisual congruency effect, point to a low degree of automaticity. This is in line with a slow developmental trajectory for automatic letter-speech integration that extends beyond the first years of reading instruction (Froyen et al., 2009).

While the superior temporal cortex became progressively more sensitive to audiovisual letter congruency, other reading-related brain areas, such as the visual word form area (VWFA), did not show similar tuning. The shifted time onset between the visual and the auditory presentation might have reduced chances to observe an audiovisual congruency effect in ventral occipitotemporal areas. It is possible that, after early activation during the visual presentation, there was no additional VWFA recruitment with spoken syllables. More research on simultaneous and non-simultaneous audiovisual presentations is needed to clarify this point. The lack of occipitotemporal effects might also relate to levels of reading automaticity, with the VWFA becoming more responsive to auditory/audiovisual stimuli as reading automaticity increases (Monzalvo, & Dehaene-Lambertz, 2013; Yoncheva, Zevin, Maurer, & McCandliss, 2010). The present findings suggest that at low levels of automaticity the left superior temporal cortex plays a crucial role in establishing cross-modal correspondences between letters and speech sounds. The VWFA does not seem to be as crucial at this stage but might become more relevant after several years of reading instruction (Froyen et al., 2009). These findings are in line with the idea that entrenched audiovisual brain networks represent an essential prerequisite for reading development that precedes the functional tuning of the VWFA (Blomert, 2011).

Previous research has reported lack of occipitotemporal response during audiovisual processing (Kapiridis et al., 2018; van Atteveldt et al., 2004), leading to the general claim that audiovisual congruency effects are more often observed in auditory than visual areas (Blomert & Froyen, 2010; van Atteveldt et al., 2004). However, such effects differ from those associated with the neural network for audiovisual speech, which requires a stronger involvement of visual areas (Calvert et al., 1998; Calvert, 2001). The source of this discrepancy might be related to the different nature of the audiovisual associations in question. While in audiovisual speech the visual component (i.e., lip movements) occurs simultaneously with speech input across the lifespan, the associations between letters and sounds are arbitrary and do not always occur simultaneously. Thus, although there is partial recycling of brain areas naturally evolved for audiovisual analysis, letter-sound associations maintain a certain degree of specificity (Blomert & Froyen, 2010).

We also found no effects in parietal areas, such as the supramarginal and angular gyri, generally thought to be involved in access to phonological representations of text (Booth, Burman, Meyer, Gitelman, Parrish, & Mesulam, 2004; Pugh et al., 2000; Schlaggar & McCandliss, 2007). This might be due to differences in experimental design: audiovisual effects in parietal areas are more often observed in comparisons of unimodal and bimodal linguistic stimuli than in comparisons of matching and mismatching audiovisual conditions (Xu et al., 2018, 2019, 2020). These parietal areas may be more involved in audiovisual letter integration than in subsequent feedback to sensory brain areas.

Finally, although our participants were early bilinguals, the present results are compatible with those reported in monolinguals (Herdman et al. 2006; Hocking & Price 2008; Karipidis et al., 2017, 2018). In addition, both writing systems learned by the children in this study (Spanish and Basque) were highly transparent and required similar learning strategies. Greater differences have been reported for late bilinguals (Bidelman & Health, 2019a, 2019b). Additional research is needed to understand to what extent neural correlates of audiovisual analysis can be generalized to diverse linguistic profiles.

In conclusion, the present study sheds light on the developmental changes of audiovisual syllable processing. Within the first months of reading instruction children progressively set up letter-sound associations, which can be detected within the first 400 ms of bimodal presentation and recruit the left superior temporal cortex. This reading-dependent brain tuning supports the idea that general mechanisms of audiovisual processing are applied (at least partially) to new arbitrary correspondences between letters and speech sounds.

## Acknowledgements

This project received funding from the European Union’s Horizon 2020 research and innovation programme under Marie Sklodowska-Curie grant agreement No 837228 (H2020-MSCA-IF-2018-837228-ENGRAVING). It was also funded by the Spanish Ministry of Economy, Industry and Competitiveness (PSI2017-82941-P), the Basque Government through the BERC 2018-2021 program, and the Agencia Estatal de Investigación through BCBL’s Severo Ochoa excellence accreditation SEV-2015-0490.

